# Computing the Internode Certainty and related measures from partial gene trees

**DOI:** 10.1101/022053

**Authors:** Kassian Kobert, Leonidas Salichos, Antonis Rokas, Alexandros Stamatakis

## Abstract

We present, implement, and evaluate an approach to calculate the internode certainty and tree certainty on a given reference tree from a collection of partial gene trees. Previously, the calculation of these values was only possible from a collection of gene trees with exactly the same taxon set as the reference tree. An application to sets of partial gene trees requires mathematical corrections in the internode certainty and tree certainty calculations. We implement our methods in RAxML and test them on empirical data sets. These tests imply that the inclusion of partial trees does matter. However, in order to provide meaningful measurements, any data set should also include trees containing the full species set.

## 1 Introduction

### 1.1 Motivation and related work

Recently Salichos and Rokas (2013) proposed a set of novel measures for quantifying the confidence for bipartitions in a phylogenetic tree (*i.e*. a leaf-labeled tree depicting the relationships between taxa). These measures are the so-called Internode Certainty (*IC*) and Tree Certainty (*TC*), which are calculated for a specific reference tree given a collection of other trees with the exact same taxon set. The calculation of their scores was implemented (Salichos *et al*., 2014) in the phylogenetic software RAxML (Stamatakis, 2014).

The underlying idea of Internode Certainty is to assess the degree of conflict of each internal branch (*i.e*. a branch connecting two internal nodes) of a phylogenetic reference tree by calculating Shannon’s Measure of Entropy (Shannon, 1948). This score is evaluated for each bipartition in the reference tree independently. The basis for the calculations are the frequency of occurrence of this bipartition and the frequencies of occurrences of a set of conflicting bipartitions from the collection of trees. In contrast to classical scoring schemes for the branches, such as simple bipartition support or posterior probabilities, the IC score also reflects to which degree the most favored bipartition is contested.

The reference tree itself can, for example, be constructed from this tree set or can be a maximum likelihood tree for a phylogenomic alignment. The tree collection may, for example, come from running multiple phylogenetic searches on the same data set, multiple bootstrap runs (Efron *et al*., 1996; Felsenstein, 1985), or from running the analyses separately on different genes, or different subsets of the genes (as done for example in Hejnol *et al*. (2009)). While for the first two cases the assumption of having the same taxon set is reasonable, this is often not the case for different genes. For example, gene sequences may be available for different subsets of taxa simply due to sequence availability or the absence of some genes in certain species.

In this paper, we show how to compute an appropriately corrected internode certainty (*IC*) on collections of partial gene trees. When using partial bipartitions for the calculation of the *IC* and *TC* scores we need to solve two problems. First, we need to calculate their respective adjusted support (analogous to the frequency of occurrence) (Section 2.1). Unlike in the standard case, with full taxon sets, this information cannot be directly obtained. Then, we also need to identify all conflicting bipartitions (Section 2).

An alternative method for calculating these frequencies has recently been independently developed by Smith *et al*. (2015). The method developed by Smith *et al*. is similar to what we denote as *lossless support* (see Section 2.1).

### 1.2 Bipartitions, Internode Certainty and Tree Certainty

We now briefly define the concepts and notations that we will use throughout the paper. Additionally, we formally define internode certainty and tree certainty.

**Bipartition** Given a taxon set *S*, a **bipartition** *B* of *S* is defined as a tuple of taxon subsets (*X, Y*) with 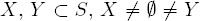 and 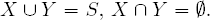. We write, *B* = *X*|*Y* = *Y*|*X*.

In phylogenetic trees, a bipartition is obtained by removing a single edge from the tree. Let *b* be an edge connecting nodes *n_1_* and *n_2_* in some unrooted phylogenetic tree *T*. The bipartition that is obtained by removing *b* is denoted by *B*(*b*), which we define as: *B*(*b*) = *X*(*n*_1_)|*X*(*n*_2_), where *X*(*n*_1_) and *X*(*n*_2_) are all taxa that are still connected to nodes *n*_1_ and *n*_2_ respectively, if branch *b* is removed.

**Trivial bipartition** We call a bipartition *B* = *X*|*Y* **trivial** if |*X*| = 1 or |*Y*| = 1.

Trivial bipartitions are uninformative, since having only a single taxon in either *X* or *Y* means that this taxon is connected to the rest of the tree. This is trivially given for any tree containing this taxon.

Bipartitions with |*X*| ≥ 2 and |*Y*| ≥ 2 are called **non-trivial**. In contrast to trivial bipartitions, non-trivial bipartitions contain information about the structure of the underlying topology. Henceforth, the term bipartition will always refer to a non-trivial bipartition.

**Sub-bipartition, super-bipartition** We denote *B*_1_ = *X*_1_|*Y*_1_ as a **sub-bipartition** of *B*_2_ = *X*_2_|*Y*_2_ if *X*_1_ ⊆ *X*_2_ and *Y*_1_ ⊆ *Y*_2_, or *X*_1_ ⊆ *Y*_2_ and *Y*_1_ ⊆ *X*_2_. The bipartition *B*_2_ is then said to be a **super-bipartition** of *B*_1_.

We also need a notion of compatibility and conflict between bipartitions.

**Conflicting bipartitions** Two bipartitions *B*_1_ = *X*_1_|*Y*_1_ and *B*_2_ = *X*_2_*Y*_2_ are **conflicting/incompatible** if there exi|s no single tree topology that explains/contains both bipartitions. Otherwise, if such a tree exists, they must be compatible. More formally, the bipartitions *B*_1_ and *B*_2_ are incompatible *if and only if* all of the following properties hold (see for example Bryant (2003)):

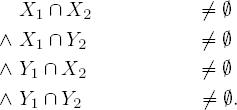

This definition of conflict and compatibility is valid irrespective of whether the taxon sets of *B*_1_ and *B*_2_ are identical or not.

**Relative frequency** Let *B*(*b*) be the bipartition induced by removing branch *b*, and let *B*^*^ be the bipartition from the tree collection that has the highest frequency of occurrence and is incompatible with *B*(*b*). Let the term *X* be the relative frequencies of the involved bipartitions. More formally, we define *X*_B_(*b*) as,

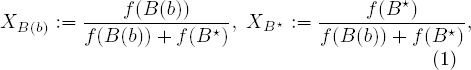

where *f* simply denotes the frequency of occurrence of a bipartition in the tree set.

For the standard case of *IC* calculations (without partial gene trees), the frequency of occurrence *f* is simply the number of observed bipartitions in the tree set. In Section 2.1 we will show how to calculate the support (adjusted frequencies) for bipartitions from partial gene trees. We compute this support using the observed frequencies of occurrence. The support for partial bipartitions can then be used analogously to the frequency of occurrence in Equation 1 for calculating the relative frequencies.

**Internode certainty** The **Internode certainty** (*IC*) score (as defined in Salichos and Rokas (2013)) is calculated using Shannon’s measure of entropy (Shannon, 1948). For a branch *b* we define *IC*(*b*) as follows:

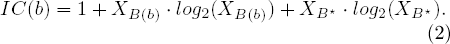

Similarly to the *IC* score, Salichos *et al*. (2014) also introduced the *ICA* (**internode certainty all**) value for each branch. However, before we formally define the *ICA* value, we need to provide some additional definitions and make some observations.

**Conflicting set** Let the set *C*^*^(*b*), as defined in Salichos *et al*. (2014), be *B*(*b*) union the set of bipartitions that conflict with *B*(*b*) and with each other, while the sum of support for elements in *C*^*^(*b*) is maximized.

In practice, the set *C*^*^(*b*) is not easy to obtain. In fact, as we show in the following observation, maximizing the sum of supports for elements in *C*^*^(*b*) renders the search for an optimal choice of *C*^*^(*b*) *NP* - *hard*.

**Observation:** Finding the optimal set *C*^*^(*b*) is *NP* - *hard*.

This can easily be seen by considering the related, known to be NP-hard, maximum weight independent set problem (Garey and Johnson, 1990). Alternatively, the similarity to the problem of constructing the asymmetric median tree, which is also known to be *NP* - *hard* (Phillips and Warnow, 1996), can be observed.

For the maximum weight independent set problem, we are confronted with an undirected graph whose nodes have weights. The task is then to find a set of nodes that maximize the sum of weights, such that no two nodes in this set are connected via an edge. A reduction from this problem to finding *C*^*^(*b*) is straight-forward. Let (*W, E*) be an undirected graph with weighted nodes *W* and edges *E*. Let *B*(*b*) = *xy*|*vz*. First, we introduce one bipartition *xz*|*vy* for every node in *W*, with support equal to the node weight. Then, for every pair of bipartitions where the corresponding nodes in *W* do not share an edge in *E*, we add four taxa that are unique to those bipartitions in such a way that they can never be compatible (consider … *ab*|*cd*… and … *ac*|*bd*…). If we find *C*^*^(*b*) for the newly introduced bipartitions, the corresponding nodes yield a maximum weight independent set.

For this reason, the definition of the *ICA*, used and implemented in Salichos *et al*. (2014), which we also use here, does not actually use *C*^*^(*b*) itself, but an approximation thereof. The set *C*(*b*) is constructed via a greedy addition strategy to approximate *C*^*^(*b*). Note that *C*(*b*) has a slightly different definition in Salichos and Rokas (2013).

Additionally, Salichos and Rokas (2013) advocate to use a threshold of 5% support frequency for conflicting bipartitions in *C*(*b*). Specifically, *C*(*b*) may only take elements 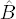 that have support

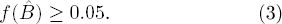

This is done to speed up the calculation. Under this restriction, the problem of maximizing the support for *C*(*b*) is no longer *NP* - *hard*. However, the search space is still large enough to warrant a greedy addition strategy instead of searching for the best solution exhaustively.

Again, let *X* denote the relative support of the bipartitions in *C*(*b*). That is,

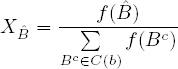

for all involved bipartitions 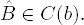.

**Internode certainty all** We can now define the *ICA* for some branch *b* as

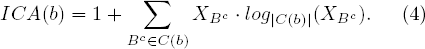

Note that *ICA*(*b*) depends on *C*(*b*). Thus, the definition for *ICA* presented here is also only an approximation. Different heuristics for constructing C(b) will yield different values for *ICA*(*b*).

Further note that iff *B*(*b*) does not have the largest frequency among all bipartitions in *C*(*b*), the *IC*(*B*) and *ICA*(*b*) scores are multiplied with —1 to indicate this. This distinction is necessary since we may have 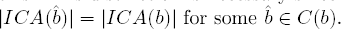 for some 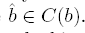. So an artificial negative value denotes that the bipartition in the reference tree is not only strongly contested, but not even the bipartition with the highest support. This can for example occur when the reference tree is the maximum-likelihood tree and the tree set contains bootstrap replicates.

From the *IC* scores and *ICA* scores the respective Tree Certainties *TC* and *TCA* can be computed. These are defined as follows:

**Tree certainty** The *TC* (**tree certainty**) and *TCA* (*tree certainty all*) scores are simply the sum over all respective *IC* or *ICA* scores, as defined in the following:

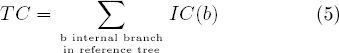

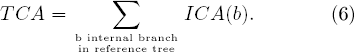

Furthermore, the *relative TC* and *TCA* scores are defined as the respective values normalized by the number of branches *b* for which *B*(*b*) is a non-trivial bipartition.

As we can see, all we need to calculate the *IC, TC, ICA*, and *TCA* scores is to calculate 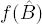 (Section 2.1) and *C*(*b*) (Section 2.2).

## 2 New Approaches: Adjusting the Internode Certainty

Now we must consider how to obtain the relevant information, namely the sets *C* and corrected support *f*, from partial bipartitions.

First, we formally define the input. We are given a so-called reference tree *T* with taxon set *S*(*T*) node set *V*(*T*) ⊇ *S*(*T*) and a set of branches *E*(*T*) ⊂ *V*(*T*) × *V*(*T*) connecting the nodes of *V*(*T*). Let 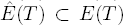 be the set of internal branches *b* for which the bipartition *B*(*b*) is non-trivial. Additionally, we are given a collection of trees 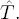. From this collection we can easily extract the set of all non-trivial bipartitions *Bip*. The bipartitions in *Bip* are used to adjust the frequency of other bipartitions. The taxon sets of the bipartitions in *Bip* are subsets of, or equal to, *S*(*T*). We call a bipartition with fewer than |*S*(*T*)| taxa a partial bipartition. A bipartition that includes all taxa from *S*(*T*) is called comprehensive or full bipartition. Similarly, a tree containing only full bipartitions is called comprehensive. From *Bip* and the bipartitions in the reference tree we can construct a set of maximal bipartitions *P* for which we will adjust the score. Bipartitions in *P* are all those bipartitions in *Bip* and the reference tree that are not sub-bipartitions of any other bipartition. We do this step, since any information contained in a sub-bipartition is also contained in the super-bipartition. Specifically, the implied gene tree (or species tree) for the super-bipartition can also explain the gene tree for all taxa in the sub-bipartition. How the frequency of occurrence of the sub-bipartition affects the frequency of occurrence of the super-bipartition is the focus of Section 2.1.

We implicitly assume that each bipartition in *P* should actually contain all taxa from *S*(*T*). To achieve this, we keep the placement of the missing taxa ambiguous. For this, we assume that each missing taxon has a uniform probability to fall into either side of the bipartition. Figure 1 gives an overview of the steps explained in the following sections.

**Figure 1:**
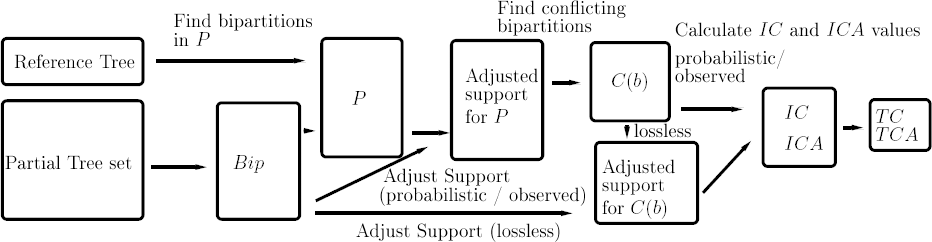
Overview of the proposed methods.

### 2.1 Correcting the Support

We aim to measure the support the given set of partial trees 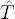 (or bipartition set *Bip*) induces for any of the bipartitions in *P*. We call this the **adjusted frequency** or **adjusted support**. If *Bip* and *P* only contain comprehensive bipartitions, the support for any given bipartition is simply equal to its frequency of occurrence.

In case of partial bipartitions, some thought must be given to the process. Imagine a comprehensive bipartition *B* = *X*|*Y* in *P* and a sub-bipartition *D* of *B* in *Bip*. Even though *D* does not exactly match *B*, it also does not contradict it. More so, it supports the super-bipartition by agreeing on a common sub-topology.

We distinguish whether the observed sub-bipartition *D* from *Bip* is allowed to support any possible bipartition, even those not observed in *Bip* and *P*, or just those we observe in *P*. There seems to be no clear answer as to which of these assumptions is more realistic. The choice is thus merely a matter of definition.

#### Support of all possible bipartitions: bilistic Support

If we assume that an observed sub-bipartition from *Bip* supports all possible super-bipartitions, not just those in *P*, with equal probability, the impact on the adjusted support of each such super-bipartition from *P*(*C*(*b*)) quickly becomes negligible. Consider the following example:

Let *B* = *X*|*Y* ∈ *P* be a super-bipartition of *D* = *x*|*y* ∈ *Bip* with |*X*\*x*| + |*Y*\*y*| = *k*. This means that *B* contains *k* taxa that *D* does not contain. There are 2^*k*^ distinct bipartitions with taxon set *X* U *Y* that also contain the constraints set by *D*. For *k* = 10 we already obtain 2^10^ = 1024 such bipartitions. Thus, the support of *D* will only increase (adjust) the support of *B* by less than one permille. More formally, let *R_B_* be the set of sub-partitions in *Bip* of the comprehensive bipartition *B* in *P* and *f_D_* the support for a partial bipartition *D* in *Bip*. Then the adjusted support for *B*, *f_B_* is

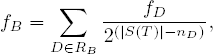

where *n_D_* is the number of taxa in *D*, and |*S*(*T*)| the number of taxa in the reference tree. We use |*S*(*T*)| in this formula, since any bipartition in *P* is is implicitly a comprehensive bipartition. By this we mean that even though we do not explicitly assign the remaining taxa from a partial bipartition *B* = *X*|*Y* in *P* to *X* or *Y*, they must belong to one of the these sets. Thus, the missing taxa in *D* have 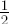 probability to belong to the same set (*X* or *Y*) each.

The effect of such an adjustment scheme is that partial bipartitions in *Bip* with fewer taxa affect the *TC* and *IC* scores substantially less than bipartitions with more taxa. This can also be observed in our computational results in Section 3. Since *f_B_*is the sum over the observed frequency times the probability of constructing the actual bipartition implied by *B*, we call this the *probabilistic* adjustment scheme.

The motivation behind the probabilistic adjustment scheme is that a partial bipartition can stem from any full bipartition that complies with the constraints induced by this partial bipartition. Furthermore, a frequency *f* > 1 for a partial bipartition can emerge due to the existence of several different, implied full bipartitions. Consider the following example: let *B*_1_ = *ABY*|*XCD* and *B*_2_ = *ABX*|*YCD* be two bipartitions from two distinct gene trees. Now, assume that taxa *X* and *Y* are not present in these gene trees (e.g., due to incomplete species sampling). In this case, the respective trees of these two gene trees only contain the same partial bipartition *B_p_* = *AB*|*CD*.

By re-distributing the frequency of *B_p_* via the probabilistic adjustment scheme to all possible bipartitions, we distribute the corresponding support among *B*_1_ and *B*_2_, as well as *B*_3_ = *ABXY*|*CD* and *B*_4_ = *AB*|*XYCD*.

#### Support of observed bipartitions: Observed Support

Now suppose that *B*_1_ and *B*_2_ are in *P* since they are present in some comprehensive or partial gene trees. Further, suppose that the bipartitions *B*_3_ and *B*_4_ (as defined above) are not in *P* since they were never observed in the tree set. Due to missing data, other partial gene trees may produce bipartition *B*_p_. In the above example for the probabilistic support, the support of *B*_p_ is not only distributed solely among *B*_1_ and *B*_2_, but also among *B*_3_ and *B*_4_, even though B3 and B4 were not observed in the tree set.

Thus, if we do not want to discard some of the frequency of occurrence when calculating the adjusted support from partial bipartitions, we can distribute their frequency of occurrence uniformly among comprehensive bipartitions in *P*. When we assume the prior distribution of bipartitions in *P* to be uniform, this process is simple. For a given partial bipartition *D* in *Bip*, with support *fD*, let *S_D_* be the set of bipartitions in *P* that are super-bipartitions of *D*. Then, *D* contributes 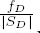 support to any *B* ∈ *S_D_* In other words, the adjusted support for each full bipartition *B* is

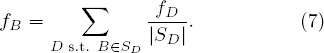

Since this distribution scheme distributes the support for each sub-bipartition among bipartitions that we observed in the tree set only, we call this the *observed* support distribution scheme.

#### Support of conflicting bipartitions: Lossless Support

One problem with the adjustment strategy explained above is that trees with more taxa typically have more bipartitions in *P* than trees with fewer taxa. For an intuitive understanding of why this can be problematic, consider the following example (also illustrated in Figure 2). Let reference bipartitions be 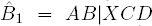 and 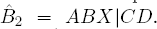 Further, let 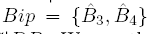 with 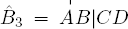 and 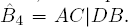 We see that 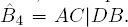 is the only, and exclusive, sub-bipartition of 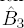 and 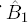 in *Bip*. Further, bipartition 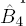 conflicts with both reference bipartitions, and no other bipartition is a super-bipartition of it. Let the bipartitions 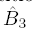 and 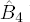 both have a frequency of occurrence of f. If we apply the above distribution scheme, bipartitions 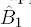 and 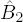 have an adjusted frequency of *f*/2, while 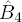 has an adjusted frequency of *f*. However, penalizing bipartitions from trees with larger taxon sets seems unwarranted. Thus, we propose a correction method that takes this into account. In order to circumvent this behavior, we choose to distribute the frequency of any sub-bipartition only to a set of conflicting super-bipartitions (namely bipartitions in *C*(*b*)). We get the following formula for the adjusted frequencies:

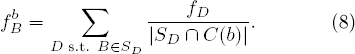

**Figure 2:**
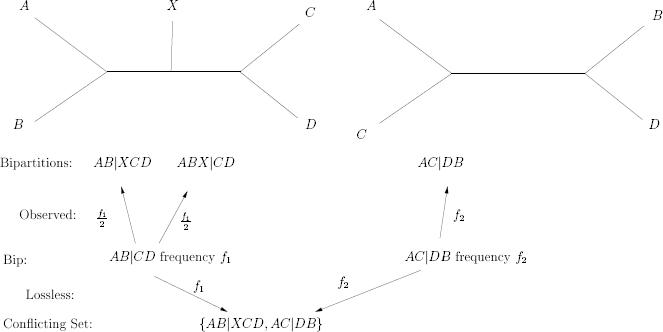
Distribution of adjusted support for observed and lossless adjustment scheme.

Where *S_D_* is defined as before. Note that the adjusted support now depends on the set of conflicting bipartitions *C*(*b*), which is defined by a branch *b*. This means that the adjusted support for a given (conflicting) bipartition must be calculated separately for each reference bipartition *B*(*b*).

This distribution scheme allocates the entire frequency of sub-bipartitions exclusively to these conflicting bipartitions. Thus, the sum of adjusted frequencies for all conflicting bipartitions is exactly equal to the sum of frequencies of occurrence of the found sub-bipartitions. For this reason, we call this the *lossless* adjustment scheme.

Note that *C*(*b*) is obtained via a greedy addition strategy depending on the adjusted support of bipartitions. Since the adjusted support according to the lossless adjustment scheme depends on *C*(*b*), we obtain a recursive definition. To alleviate this, we simply precompute the above explained probabilistic adjustment scheme to obtain an adjusted support for each bipartition. The set of conflicting bipartitions *C*(*b*) is then found with respect to the probabilistically adjusted support values. Then, using *C*(*b*), the actual lossless support adjustment is calculated and replaces the probabilistic support in the calculation of *IC* and *ICA* values.

For the above example we get the following. Let 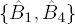 be the set of conflicting bipartitions. Then, the support for 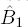 and 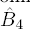 after applying the lossless distribution scheme is *f* for both bipartitions, which is the desired behavior for this distribution scheme.

### 2.2 Finding Conflicting Bipartitions

To construct *C*(*b*) greedily, as proposed above, the support of the bipartitions must be known. However, the lossless support adjustment scheme explained above is only reasonable on a set of conflicting bipartitions (for example, *C*(*b*) itself). To avoid this recursive dependency, we first compute an adjusted support that does not depend on *C*(*b*) for this case. (Here we use the probabilistic adjusted support, as explained in Section 2.1, to obtain an initial adjusted support.) Then, a greedy algorithm is used to approximate the set *C*(*b*) with the highest sum of adjusted support with respect to the initial adjustment. Once *C*(*b*) is obtained, the support for all bipartitions in *C*(*b*) is adjusted using the new method, which depends on a set of conflicting bipartitions. These new values then replace the initial estimate via the first adjustment scheme.

Keeping the above in mind, we can easily construct *C*(*b*) from *P* for every branch *b* in *E*(*T*). Note that we also defined the reference bipartition *B*(*b*) to be in *C*(*b*). Thus, we simply start with *B*(*b*) and iterate through the elements of *P* in decreasing order of adjusted support (if we are to apply the probabilistic or lossless distribution scheme, the probabilistic adjusted support is used in this step. Similarly, the observed adjusted support is used, if this distribution is desired) and add every bipartition that conflicts with all other bipartitions added to *C*(*b*) so far. During this process the threshold given in Equation 3 is applied.

Given *B*(*b*), *C*(*b*), and *Bip* we can calculate the *IC* and *ICA* values as defined in Equations 2 and 4 under the *probabilistic* or *observed* adjustment schemes. For the *lossless* adjustment scheme, the actual adjusted frequencies have to be calculated separately for each bipartition in C(b) for all reference bipartitions b in this step.

### 2.3 Example

We now present a simple example for calculating the IC score under the different adjustment schemes. To this end, we analyze the tree set shown in Figure 3. From these trees we initially extract the following bipartition lists:

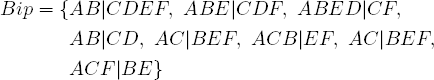

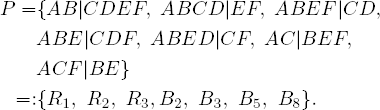

We can now immediately calculate the probabilistic and observed support for bipartitions in *P*. As mentioned before, the lossless adjustment can only be calculated on sets of conflicting bipartitions. Let 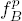 and 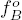 be the probabilistic and observed support of a bipartition *B*. Further, let 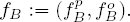

Then, as *B*_1_ in the Figure is exactly identical to *R*_1_, and *B*_4_ is a sub-bipartition of *R*_1_ with 2 missing taxa, 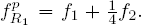 At the same time, *R*_1_ is the only super-bipartition of *B*_1_. However, two other bipartitions, namely *R*_3_ and *B*_2_, are super-bipartitions of *B*_4_. Thus, we obtain 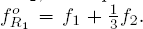 All other bipartitions in *P* can be scored analogously to obtain the following probabilistic and observed support value pairs:

**Figure 3:**
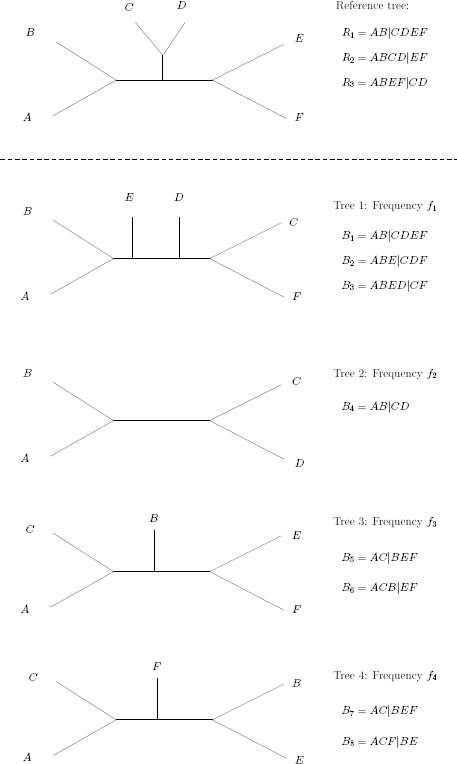
Example tree set for *IC* calculations.

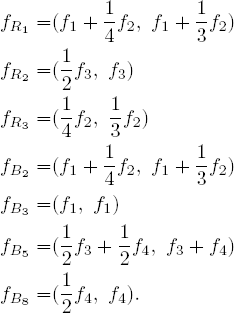

Given the above, we can now calculate the *IC* scores for bipartitions *R*_1_, *R*_2_, and *R*_3_. Assume that we have the following frequencies, *f*_1_ = 3, *f*_2_ = 4, *f*_3_ = 6, and *f*_4_ = 6. Bipartition *R*_1_ = *AB*|*CDEF* conflicts with both, *B*_5_ = *AC*|*BEF*, and *B*_8_ = *ACF*|*BE*. However, since *B*_5_ and *B*_8_ do not conflict with each other, only one of them is included in the list of conflicting bipartitions. Since *B*_5_ has a higher adjusted support than *B*_8_, we include *B*_5_. If *b* is the branch that gives rise to bipartition *R*_1_ in the reference tree, then *C*(*b*) = {*R*_1_,*B*_5_}. Under the probabilistic adjustment scheme we obtain:

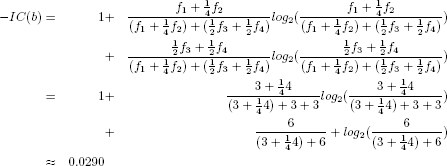

The negative value of *IC*(*b*) is due to the fact that, under the observed adjustment scheme, *B*_5_ has a higher adjusted support than *R*_1._ Similarly, under the observed adjustment scheme we obtain:

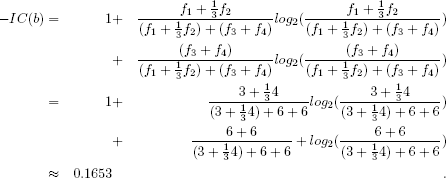

Given *C*(*b*), we can now also compute the lossless adjusted support. We obtain a support of *f*_1_ + *f*_2_ = 7 for *R*_1_, and a support of *f*_3_ + *f*_4_ = 6 + 6 for *B*_5_. With these numbers at hand, we can calculate the IC score under lossless adjustment as:

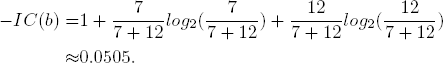

This can be done analogously for bipartitions *R*_2_ and *R*_3_. For *R*_2_ = *ABCD*|*EF* we observe three conflicting bipartitions: *B*_2_ = *ABE*|*DCF*, *B*_3_ = *ABED*|*CF*, and *B*_8_ = *ACF*|BE. The corresponding frequencies for the above bipartitions are:

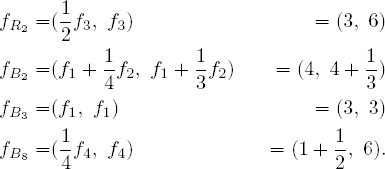

Under the probabilistic support, we thus obtain *C*(*b*) = {*R*_2_, *B*_2_}, where *b* is the branch that corresponds to the reference bipartition with *R*_2_ = *B*(*b*). However, the set of conflicting bipartitions is different for the observed adjustment scheme. Here, *C*(*b*) = {*R*_2_, *B*_8_}. As a consequence we obtain the following IC scores:

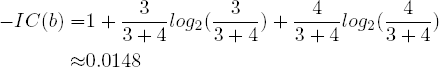

under the probabilistic scheme, and

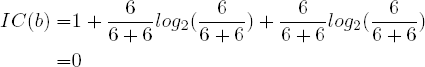

under the observed adjustment scheme. The adjusted frequencies for bipartitions *R*_2_ and *B*_2_, under the lossless adjustment scheme, are *f*_3_ = 6 and *f*_1_ + *f*_2_ = 7, respectively. Thus, the *IC* score is

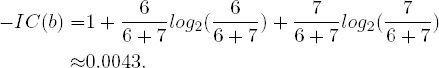

For reference bipartition *R*_3_ = *ABEF*|*CD*, there is only one conflicting bipartition in *P*, namely *B*_3_ = *ABED*|*CF*. Thus, the calculation of *IC*(*b*) is straight-forward (as before *b* is the branch inducing the reference bipartition: *R*_3_). Under the probabilistic scheme we obtain:

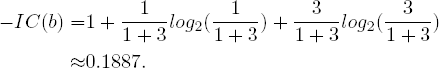

Under the observed adjustment we get:

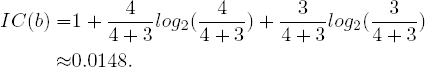

Finally, under the lossless adjustment scheme we obtain:

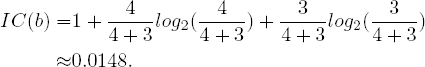

## 3 Results and Discussion

For implementing the methods described in Section 2, we used the framework of the RAxML (Stamatakis, 2014) software (version 8.1.20).

The resulting proof of concept implementations and all data sets used for our experiments in Sections 3.1 and 3.2 (as well as the above example of Section 2.3) are available at https://github.com/Kobert/ICTC. Usage of the software is explained there as well. The probabilistic and lossless distribution schemes are also included in the latest production level version of RAxML (https://github.com/stamatak/standard-RAxML, version 8.2.4).

We chose to omit the implementation for the observed support adjustment from the official RAxML release, as it does not seem to offer any advantages over the other two methods.

### 3.1 Accuracy of the Methods

In this section we asses the accuracy of the proposed adjustment schemes. For this reason, we re-analyze the yeast data set originally presented in Salichos and Rokas (2013). The comprehensive trees in the data set contain 23 taxa. After applying some filtering techniques to the genes, we obtained a set of 1275 gene trees. In the filtering step, genes are discarded if (i) the average sequence length is less than 150 characters, or (ii) more than half the sites contain indels after alignment. In Salichos and Rokas (2013), a slightly smaller subset of 1070 trees is used.

To understand which adjustment scheme better recovers the underlying truth, we randomly prune taxa from this comprehensive tree set and compare the results between adjustment schemes. Evidently, a “good” adjustment scheme will yield *IC* and *ICA* values that are as similar as possible to the *IC /ICA* values of the comprehensive tree set. Thus we consider the *IC/ICA* on the comprehensive tree set as the correct values.

For each of the 1275 trees, we select and prune a random number of taxa. We draw the numbers of taxa to prune per tree from a geometric distribution with parameter *p*. We use a geometric distribution because the expectation is that thereby we will retain *p*. 1275 comprehensive trees, for which 0 taxa have been pruned. An additional restriction is that each pruned tree must comprise at least 4 taxa to comprise at least one non-trivial bipartition. Given the number of taxa we wish to prune, we select taxa to prune uniformly at random using the *newick-tools* toolkit^1^.

Using different values for *p* we generate four partial tree sets. For each of these tree sets, we conduct analyses including all 1275 trees (comprehensive and partial). We compare the results to the *IC/ICA* scores for 1275 comprehensive trees.

Similarly, in a second round of experiments we compare the results obtained by removing *all* comprehensive trees from the tree sets to the reference *IC* and *ICA* scores for the comprehensive tree set.

To quantify which correction method yields more accurate results, we define the following distance/accuracy measure. Let *IC*(*b*) be the inter node certainty for branch *b* if no taxa are pruned. Similarly, let *IC^A^*(*b*) be the internode certainty for the same branch *b* under an adjustment scheme for a data set *with* partial gene trees. The accuracy *D* of an adjustment scheme is then defined as:

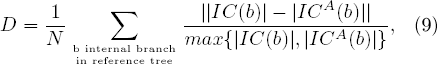

where *N* is the number of internal branches in the reference tree (*N* = 20 for our test data set). The measure *D* is the average, weighted, component-wise difference between the two results. A low value of *D* indicates high similarity between the results. Furthermore, by definition, *D* ranges between 0 and 1.

Table 1 depicts this distance *D* for the different tree sets and adjustment schemes we tested. As we can see, the probabilistic and observed adjustment methods are more accurate than the lossless method.

In Table 2 we observe that the probabilistic and observed adjustment schemes are not more accurate than the lossless method for tree sets that only contain partial gene trees. From Table 3 it also becomes evident that the lossless adjustment scheme tends to overestimate the *IC* and *ICA* values less frequently than the two alternative methods.

Another important observation is that, in most cases, accuracy decreases for any adjustment scheme when analyzing tree sets that exclusively contain partial gene trees. Intuitively, this can be explained by the fact that (i) we have less trees to base our analysis on, and (ii) only the reference bipartitions now contain all 23 taxa. Since a partial bipartition distributes its frequency among all its super-bipartitions in *P*, it is intuitively clear that bipartitions with more taxa are more likely to accumulate distributed frequencies from more sub-bipartitions than bipartitions with fewer taxa. Conflicting bipartitions (with less than 23 taxa) are thus not assigned sufficient support to compete with the reference bipartitions. This behavior can be observed in Table 3. There, we display the numbers of times the certainty in a branch under the different adjustment schemes was higher than the certainty obtained from the comprehensive trees.

### 3.2 Empirical Data Analyses

In this section we present an additional, yet different, analysis of the above yeast data set. We do not only use the 1275 comprehensive trees, but now also include additional partial gene trees. After applying the aforementioned filters again (3.1), the tree set comprises 2494 trees. The comprehensive trees are the same 1275 trees as in Section 3.1. The remaining 1219 trees are partial trees. The number of taxa in these partial trees ranges from 4 to 22 (see Figure 4 for the exact distribution of taxon numbers over partial gene trees). Unlike in Section 3.1, these partial trees are not simulated, but the result of phylogenetic analyses on the corresponding gene alignments.

In addition, we also analyze a gene tree set from avian genomes. The data was previously published in Jarvis *et al*. (2015). Here, we analyze a subset of 2000 gene trees with up to 48 taxa. Of these trees, 500 contain the full 48 taxa while the remaining trees contain either 47 taxa (500 trees) or 41-43 taxa (1000 trees). The taxon number distribution over trees is provided in Figure 5.

First, we report the results for the yeast data set. We present the IC and ICA scores for all internal branches under the three adjustment schemes and compare them to the scores obtained for the subset of comprehensive trees. Figure 6 shows the topology of the reference tree. Tables 4 and Table 5 show the respective *IC* and *ICA* values.

The values for the individual *IC* and *ICA* scores *can* be higher for the lossless adjustment scheme than for the probabilistic adjustment scheme and the observed adjustment scheme. However, the relative *TC* and *TCA* values suggest that the lossless adjustment attributes a lower certainty to individual bipartitions as well as the entire tree. The actual values are 0.298 for the relative *TC* score and 0.322 for the relative *TCA* score for the lossless adjustment; 0.389 and 0.399 for the probabilistic adjustment; and 0.339 and 0.364 for the observed adjustment scheme.

**Table 1:** Differences *D* in *IC/ICA* scores, between the scores calculated by the adjustment schemes and the reference scores for the comprehensive tree set.

|  | *IC* |  |  |  | *ICA* |  |  |  |
| --- | --- | --- | --- | --- | --- | --- | --- | --- |
| | $p = 0.1$ | $p = 0.3$ | $p = 0.5$ | $p = 0.7$ | $p = 0.1$ | $p = 0.3$ | $p = 0.5$ | $p = 0.7$ |
| Probabilistic | 0.31 | 0.20 | 0.18 | 0.08 | 0.26 | 0.18 | 0.18 | 0.12 |
| Observed | 0.42 | 0.27 | 0.15 | 0.07 | 0.39 | 0.25 | 0.19 | 0.08 |
| Lossless | 0.65 | 0.44 | 0.24 | 0.17 | 0.60 | 0.44 | 0.28 | 0.15 |

**Table 2:**
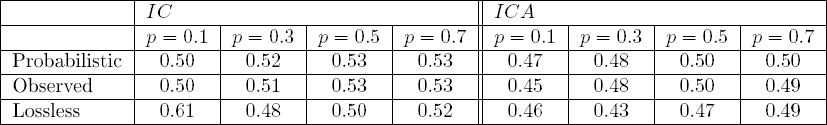
Differences *D* in *IC /ICA* scores, between the pruned tree sets *only* containing partial gene trees and the reference values.

|  | *IC* |  |  |  | *ICA* |  |  |  |
| --- | --- | --- | --- | --- | --- | --- | --- | --- |
| | $p = 0.1$ | $p = 0.3$ | $p = 0.5$ | $p = 0.7$ | $p = 0.1$ | $p = 0.3$ | $p = 0.5$ | $p = 0.7$ |
| Probabilistic | 0.50 | 0.52 | 0.53 | 0.53 | 0.47 | 0.48 | 0.50 | 0.50 |
| Observed | 0.50 | 0.51 | 0.53 | 0.53 | 0.45 | 0.48 | 0.50 | 0.49 |
| Lossless | 0.61 | 0.48 | 0.50 | 0.52 | 0.46 | 0.43 | 0.47 | 0.49 |

**Table 3:** Fraction of branches for which the adjusted *IC /ICA* scores are higher than the *IC /ICA* reference scores. The top table contains values for all three adjustment schemes if all trees (comprehensive and simulated partial) are included in the analysis. The bottom table shows the values for all three methods if *only* partial trees are analyzed.

|  | *IC* |  |  |  | *ICA* |  |  |  |
| --- | --- | --- | --- | --- | --- | --- | --- | --- |
| | $p = 0.1$ | $p = 0.3$ | $p = 0.5$ | $p = 0.7$ | $p = 0.1$ | $p = 0.3$ | $p = 0.5$ | $p = 0.7$ |
| All trees |  |  |  |  |  |  |  |  |
| Probabilistic | 0.4 | 0.35 | 0.35 | 0.15 | 0.25 | 0.25 | 0.2 | 0.15 |
| Observed | 0.15 | 0.3 | 0.4 | 0.2 | 0.2 | 0.2 | 0.2 | 0.1 |
| Lossless | 0.1 | 0.25 | 0.15 | 0.25 | 0.2 | 0.2 | 0.25 | 0.1 |
| Partial trees |  |  |  |  |  |  |  |  |
| Probabilistic | 0.8 | 0.8 | 0.85 | 0.85 | 0.8 | 0.8 | 0.85 | 0.85 |
| Observed | 0.65 | 0.75 | 0.8 | 0.85 | 0.65 | 0.75 | 0.8 | 0.85 |
| Lossless | 0.3 | 0.65 | 0.75 | 0.8 | 0.25 | 0.65 | 0.75 | 0.8 |

**Table 4:**
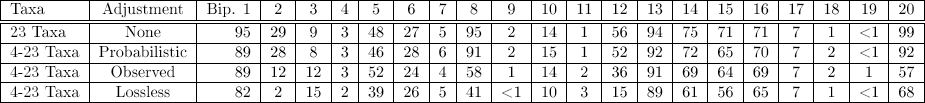
IC scores for all non-trivial bipartitions multiplied by 100 and rounded down. The bipartition labels are shown in Figure 6. The data set can either consist of only full trees (23 taxa), or partial and full trees (4-23 taxa).

**Figure 6:**
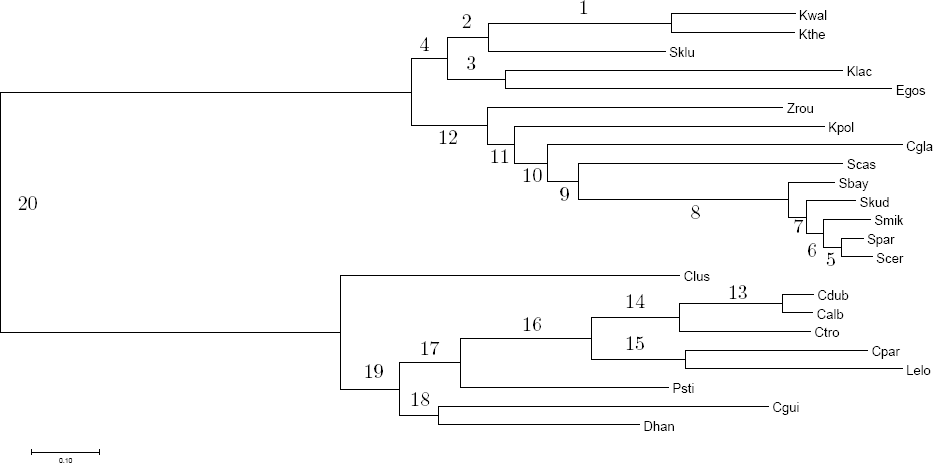
Bipartition numbers corresponding to the presented tables, for the yeast data set. Taxon key: Kwal: *Kluyveromyces waltii*, Kthe: *Kluyveromyces thermotolerans*, Sklu: *Saccharomyces kluyveri*, Klac: *Kluyveromyces lactis*, Egos: *Eremothecium gossypii*, Zrou: *Zygosacharomyces rouxii*, Kpol: *Kluyveromyces polysporus*, Cgla: *Candida glabrata*, Seas: *Saccharomyces castella*, Sbay: *Saccharomyces bayanus*, Skud: *Saccharomyces kudriavzevii*, Smik: *Saccharomyces mikatae*, Spar: *Saccharomyces paradoxus*, Scer: *Saccharomyces cerevisiae*, Clus: *Candida lusitaniae*, Cdub: *Candida dubliniensis*, Calb: *Candida albicans*, Ctro: *Candida tropicalis*, Cpar: *Candida parapsilosis*, Lelo: *Lodderomyces elongisporus*, Psti: *Pichia stipitis*, Cgui: *Candida guilliermondii*, Dhan: *Debaryomyces hansenii*

**Table 5:** ICA scores for all non-trivial bipartitions multiplied by 100 and rounded down. The bipartition labels are shown in Figure 6. The data sets again either consist of only full trees (23 taxa), or partial and full trees (4-23 taxa).

| Taxa | Adjustment | Bip. 1 | 2 | 3 | 4 | 5 | 6 | 7 | 8 | 9 | 10 | 11 | 12 | 13 | 14 | 15 | 16 | 17 | 18 | 19 | 20 |
| --- | --- | --- | --- | --- | --- | --- | --- | --- | --- | --- | --- | --- | --- | --- | --- | --- | --- | --- | --- | --- | --- |
| 23 Taxa | None | 95 | 23 | 7 | 8 | 48 | 25 | 14 | 95 | 3 | 12 | 2 | 45 | 94 | 75 | 71 | 71 | 7 | 8 | 9 | 98 |
| 4-23 Taxa | Probabilistic | 89 | 21 | 6 | 13 | 46 | 26 | 14 | 91 | 3 | 11 | 1 | 38 | 92 | 72 | 60 | 70 | 25 | 7 | 11 | 92 |
| 4-23 Taxa | Observed | 89 | 15 | 9 | 12 | 52 | 24 | 12 | 58 | 2 | 11 | 11 | 34 | 91 | 69 | 59 | 69 | 24 | 7 | 11 | 57 |
| 4-23 Taxa | Lossless | 82 | 13 | 10 | 7 | 39 | 27 | 13 | 46 | 3 | 9 | 8 | 29 | 89 | 61 | 49 | 65 | 7 | 5 | 5 | 68 |

By comparing the 23-taxa yeast species tree values without adjustment against the three approaches that contain both complete and missing data (probabilistic, observed and lossless), we can conclude that, overall, the values appear very similar and they tend to provide additional support for the reference topology. Among the adjustment strategies, the probabilistic adjustment yields values that are closest to those obtained by the analysis of only comprehensive trees. This is expected, since for the probabilistic adjustment, smaller bipartitions contribute less to the overall scores than larger bipartitions. Full bipartitions/trees are thus affecting the outcome most under this adjustment scheme.

**Figure 4:**
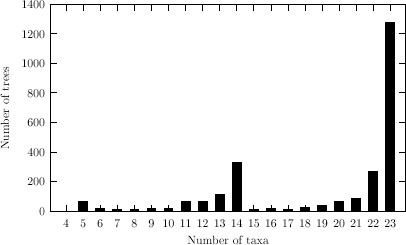
Distribution of taxon number over trees in the yeast data.

**Figure 5:**
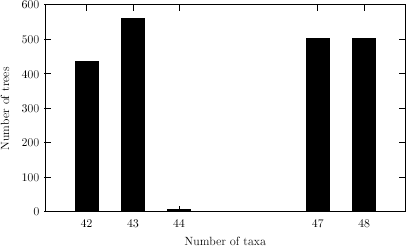
Distribution of taxon number over trees in the avian data.

Previous ambiguous bipartitions, concerning for example the placement of species like *S. castelii* (conf. bipartitions 9 and 8), *C. lusitaniae* (conf. bipartitions 20 and 19), *D. hansenii* (bipartition 18), and *K. lactis* (bipartition 3), remain equally uncertain, showing very similar (close to 0) IC and ICA values.

The split between the *Candida* and *Saccharomyces* clade (bipartition 20) is well documented in the literature (Fitzpatrick *et al*., 2006; Dujon, 2010; Salichos and Rokas, 2013). The same holds for bipartition 8, the *Saccharomyces ’sensu stricto’* clade (Rokas *et al*, 2003; Kurtzman and Robnett, 2006; Salichos and Rokas, 2013). Thus, a high certainty for these bipartitions is expected. As we can see in Table 4, the analysis of only comprehensive trees supports these two bipartitions with IC values of 0.99 for bipartition 20, and 0.95 for bipartition 8. However, the generally conservative lossless distribution approach, as well as the observed support adjustment scheme, provide reduced certainty for these two bipartitions; the divergence of *Candida* from the *Saccharomyces* clade (bipartition 20) is, for the lossless distribution scheme, depicted with an IC value of 0.68, and the *Saccharomyces ’sensu stricto’* clade (bipartition 8) obtains an IC score of 0.41; the observed adjusted support for these bipartitions is reduced to 0.57 for bipartition 20, and 0.58 for bipartition 8. The probabilistic adjusted IC values for the branches inducing these splits are 0.92 for bipartition 20, and 0.91 for bipartition 8. A similar behavior can be seen for the ICA values.

In addition, under the lossless adjustment, the previously resolved placement of *Z. rouxii* (a clade with relatively low gene support frequency of 62% in (Salichos and Rokas, 2013)) remains unresolved with IC and ICA values of 0.15 and 0.29 respectively.

Next, we analyze the behavior of the adjustment schemes if *only* partial trees are provided. See Tables 6 and 7.

The relative *TC* (and *TCA)* that result from these calculations are 0.668 (0.651) for the probabilistic distribution, 0.499 (0.532) for the observed distribution, and 0.394 (0.407) for the lossless distribution scheme. The relative *TC* and *TCA* without correction (obtained from the values shown in Tables 4 and 5), for trees with full taxon sets, are 0.406 and 0.409. The higher *TC* and *TCA* values obtained for the former two adjustment methods suggest that these approaches are not providing the conflicting bipartitions with a sufficiently adjusted support to compare to the reference bipartition. The reference bipartitions always contain 23 taxa for this data set. Now however, no conflicting bipartition can have that many taxa, as comprehensive trees are not included in the above analysis of only partial trees.

**Table 6:** IC scores for all non-trivial bipartitions multiplied by 100 and rounded down. The bipartition labels are shown in Figure 6. Here, the data set only contains trees with partial taxon sets.

| Taxa | Adjustment | Bip. 1 | 2 | 3 | 4 | 5 | 6 | 7 | 8 | 9 | 10 | 11 | 12 | 13 | 14 | 15 | 16 | 17 | 18 | 19 | 20 |
| --- | --- | --- | --- | --- | --- | --- | --- | --- | --- | --- | --- | --- | --- | --- | --- | --- | --- | --- | --- | --- | --- |
| 23 Taxa | None | 95 | 29 | 9 | 3 | 48 | 27 | 5 | 95 | 2 | 14 | 1 | 56 | 94 | 75 | 71 | 71 | 7 | 1 | <1 | 99 |
| 4-22 Taxa | Probabilistic | 93 | 64 | 61 | 58 | 72 | 66 | 59 | 85 | 39 | 46 | 43 | 64 | 95 | 77 | 83 | 78 | 56 | 49 | 47 | 93 |
| 4-22 Taxa | Observed | 89 | 23 | 58 | 36 | 80 | 75 | 70 | 80 | 1 | 1 | <1 | 20 | 93 | 79 | 82 | 78 | 54 | 13 | 16 | 43 |
| 4-22 Taxa | Lossless | 80 | 24 | 58 | 12 | 66 | 57 | 32 | 68 | 24 | 12 | 12 | 2 | 88 | 54 | 42 | 49 | 43 | 12 | 38 | 7 |

**Table 7:** ICA scores for all non-trivial bipartitions multiplied by 100 and rounded down. The bipartition labels are shown in Figure 6. Again, the data set only contains trees with partial taxon sets.

| Taxa | Adjustment | Bip. 1 | 2 | 3 | 4 | 5 | 6 | 7 | 8 | 9 | 10 | 11 | 12 | 13 | 14 | 15 | 16 | 17 | 18 | 19 | 20 |
| --- | --- | --- | --- | --- | --- | --- | --- | --- | --- | --- | --- | --- | --- | --- | --- | --- | --- | --- | --- | --- | --- |
| 23 Taxa | None | 95 | 23 | 7 | 8 | 48 | 25 | 14 | 95 | 3 | 12 | 2 | 45 | 94 | 75 | 71 | 71 | 7 | 8 | 9 | 98 |
| 4-22 Taxa | Probabilistic | 93 | 64 | 54 | 51 | 72 | 66 | 59 | 85 | 40 | 46 | 34 | 58 | 95 | 77 | 83 | 78 | 56 | 43 | 45 | 93 |
| 4-22 Taxa | Observed | 89 | 23 | 48 | 33 | 80 | 75 | 70 | 80 | 17 | 20 | 18 | 20 | 93 | 79 | 82 | 78 | 54 | 29 | 24 | 43 |
| 4-22 Taxa | Lossless | 80 | 27 | 58 | 24 | 66 | 57 | 29 | 68 | 24 | 11 | 12 | 2 | 88 | 54 | 42 | 49 | 43 | 12 | 38 | 22 |

Analyzing the second data set with a total of 2000 trees yields similar results. See Table 3.2 for the *TC* and *TCA* values for this data set. Again, the values of the analysis restricted to a comprehensive tree set are compared to the results obtained when including partial gene trees, and restricting the analysis to partial gene trees. Specifically, we see that the probabilistic support for analyzing the full data set, of 2000 trees, again gives *TC* values more closely in accordance with the values obtained for the analysis restricted to the 500 full trees than the lossless adjustment scheme.

**Table 8:** *IC* and *ICA* scores for different subsets of the data set for the probabilistic and lossless distribution schemes.

| Taxa | adjustment | $TC$ | $TCA$ |
| --- | --- | --- | --- |
| 48 taxa | None | -3.14 | -3.17 |
| 41-48 Taxa | Probabilistic | -2.44 | 7.72 |
| 41-48 Taxa | Lossless | -5.05 | -1.35 |
| 41-47 Taxa | Probabilistic | 9.34 | 15.75 |
| 41-47 Taxa | Lossless | 6.01 | 6.01 |

Here, the tree set does not support the reference tree well (as evident by the negative *TC)*. At the same time, the *TCA* under the probabilistic adjustment scheme is actually positive.

For this data set, the discrepancy can be explained by the fact that the most frequent conflicting bipartitions are not supported by much more than the second most supported conflicting bipartition. If the support for the reference bipartition is much smaller than that of the most frequent conflicting bipartition, the internode-certainty will approach — 1. Let the support for the most frequent conflicting bipartition be *f*. As the support of the second most frequent conflicting bipartition approaches *f*, the *ICA* value tends towards 0.0. If the reference bipartition is the bipartition with the highest adjusted support in *C*(*b*), this effect is less pronounced.

For the analysis of partial bipartitions only, we again see that the conflicting bipartitions are not as well supported under any tested adjustment scheme. Again, the lossless adjustment scheme yields decreased certainty. Thus, we advocate that this adjustment scheme is used if one wants to reduce the risk of overestimating certainties.

## 4 Conclusion

We have seen that the inclusion of partial trees into any certainty estimation is beneficial, as the partial trees do contain information that is not necessarily contained in the full/comprehensive trees. This is evident by the different *TC* and *TCA* scores we obtained for the empirical data sets.

Further, the selection of the most appropriate adjustment scheme depends on the data at hand. The lossless adjustment scheme is most appropriate for tree sets that do not contain any comprehensive trees, since it yields more conservative certainty estimates. For gene tree sets that contain comprehensive as well as partial trees, the probabilistic and observed adjustment schemes yield results that are more accurate with respect to the reference *IC* and *ICA* values.

In general, we advocate the inclusion of (some) comprehensive trees in any analysis that also includes partial trees. This is motivated by the fact that the pruned data sets that contained comprehensive trees generally yielded more accurate results than tree sets not containing comprehensive trees.

## Footnotes

1 https://github.com/xflouris/newick-tools

## References

Bryant, D. 2003. A classification of consensus methods for phylogenies. In M. Janowitz, F.-J. Lapointe, F. McMorris, B. Mirkin, and F. Roberts, editors, Bioconsensus, DIMACS. AMS-, pages 163-184.

Dujon, B. 2010. Yeast evolutionary genomics. Nature Reviews Genetics, 11(7): 512-524.

Efron, B., Halloran, E., and Holmes, S. 1996. Bootstrap confidence levels for phylogenetictrees. Proceedings of the. National Academy of Sciences, 93(23).

Felsenstein, J. 1985. Confidence limits on phylogenies: an approach using the bootstrap. Ann. Stat., 39: 783–791.

Fitzpatrick, D., Logue, M., Stajich, J., and Butler, G. 2006. A fungal phylogeny based on 42 complete genomes derived from supertree and combined gene analysis. BMC Evolutionary Biology, 6(1): 99.

Garey, M. R. and Johnson, D. S. 1990. Computers and Intractability; A Guide to the Theory of NP-Completeness. W. H. Freeman & Co., New York, NY, USA.

Hejnol, A., Obst, M., Stamatakis, A., Ott, M., Rouse, G. W., Edgecombe, G. D., Martinez, P., Bagunà, J., Bailly, X., Jondelius, U., Wiens, M., Miiller, W. E. G., Seaver, E., Wheeler, W. C, Martindale, M. Q., Giribet, G., and Dunn, C. W. 2009. Assessing the root of bilaterian animals with scalable phylogenomic methods. Proceedings of the Royal Society of london B: Biological Sciences.

Jarvis, E., Mirarab, S., Aberer, A., Li, B., Houde, P., Li, C, Ho, S., Faircloth, B., Nabholz, B., Howard, J., Suh, A., Weber, C, da Fonseca, R., Alfaro-Núñez, A., Narula, N., Liu, L., Burt, D., Ellegren, H., Edwards, S., Stamatakis, A., Mindell, D., Cracraft, J., Braun, E., Warnow, T., Jun, W., Gilbert, M., and Zhang, G. 2015. Phylogenomic analyses data of the avian phylogenomics project. GigaScience, 4(1).

Kurtzman, C. P. and Robnett, C. J. 2006. Phylogenetic relationships among yeasts of the ’Saccharomyces complex’ determined from multigene sequence analyses. FEMS Yeast Research, 3(4): 417–432.

Phillips, C. and Warnow, T. J. 1996. The asymmetric median tree — a new model for building consensus trees. Discrete Applied Mathematics, 71(1-3): 311 -335.

Rokas, A., Williams, B. L., King, N., and Carroll, S. B. 2003. Genome-scale approaches to resolving incongruence in molecular phylogenies. Nature, 425(6960): 798–804.

Salichos, L. and Rokas, A. 2013. Inferring ancient divergences requires genes with strong phylogenetic signals. Nature.

Salichos, L., Stamatakis, A., and Rokas, A. 2014. Novel information theory-based measures for quantifying incongruence among phylogenetic trees. Molecular Biology and Evolution.

Shannon, C. E. 1948. A mathematical theory of communication. The Bell System Technical Journal, 27.

Smith, S., Moore, M., Brown, J., and Yang, Y. 2015. Analysis of phylogenomic datasets reveals conflict, concordance, and gene duplications with examples from animals and plants. BMC Evolutionary Biology, 15(1): 150.

Stamatakis, A. 2014. Raxml version 8: A tool for phylogenetic analysis and post-analysis of large phylogenies. Bioinformatics.

